# A Framework for Fast, Large-scale, Semi-Automatic Inference of Animal Behavior from Monocular Videos

**DOI:** 10.1101/2023.07.31.551177

**Authors:** Eric Price, Pranav C. Khandelwal, Daniel I. Rubenstein, Aamir Ahmad

## Abstract

An automatic, quick, accurate, and scalable method for animal behavior inference using only videos of animals offers unprecedented opportunities to understand complex biological phenomena and answer challenging ecological questions. The advent of sophisticated machine learning techniques now allows the development and implementation of such a method. However, apart from developing a network model that infers animal behavior from video inputs, the key challenge is to obtain sufficient labeled (annotated) data to successfully train that network - a laborious task that needs to be repeated for every species and/or animal system. Here, we propose solutions for both problems, i) a novel methodology for rapidly generating large amounts of annotated data of animals from videos and ii) using it to reliably train deep neural network models to infer the different behavioral states of every animal in each frame of the video. Our method’s workflow is bootstrapped with a relatively small amount of manually-labeled video frames. We develop and implement this novel method by building upon the open-source tool Smarter-LabelMe, leveraging deep convolutional visual detection and tracking in combination with our behavior inference model to quickly produce large amounts of reliable training data. We demonstrate the effectiveness of our method on aerial videos of plains and Grévy’s Zebras (*Equus quagga* and *Equus grevyi*). We fully open-source the code^1^ of our method as well as provide large amounts of accurately-annotated video datasets^2^ of zebra behavior using our method. A video abstract of this paper is available here^3^.

## 1. Introduction

One of the cornerstones in the field of animal behavior is observing animals in the wild [1]. Meticulous field observations have led to novel insights into how animals behave at the individual and interact at the group level in the animal kingdom. For example, observations made in the wild have provided novel insights into various aspects of animal behavior research including mating systems, parental care, foraging strategies, social structures, conspecific communication, and altruistic behaviors [17, 19].

Traditionally, the quantification of animal behavior has relied on direct observations and opportunistic sampling by researchers/volunteers in the animal’s natural habitat [11]. Observers spend hours in the field, carefully documenting behavioral events and collecting data on individual animals. While this approach has provided valuable information, it is labor-intensive, time-consuming, and often limited in capturing the full complexity and subtlety of the animal’s behavior, especially over large spatiotemporal scales and group sizes.

Overcoming the issues related to manual observations has seen the rapid adoption of image/video recording devices, remote sensing techniques, and bio-loggers to quantify the behavior of animals [8, 22]. These techniques allow the collection of large datasets rich in diverse behaviors in the animal’s natural habitat [16]. Video-based approaches are especially attractive since they do not involve handling the animal and enable capturing the behavior along with its behavioral context [3]. Video-based techniques include camera traps, smartphone cameras, and more recently, unmanned aerial vehicles (UAV) that can follow animals over large spatiotemporal scales and circumvent line of sight obstructions imposed by enviromental objects [21, 5, 20, 2].

However, image/video-based approaches come with their own set of challenges. Large amounts of recorded data must be manually or automatically analyzed to extract behavior events. Manual analysis of images or videos require minimal computational know-how but are cumbersome, time-consuming, and to a large extent, subjective – the extracted data is dependent on the annotator, which can vary depending on a trained researcher versus a volunteer [3]. Automatic analysis can be significantly faster and provide standardized and objective behavior measurements, albeit if implemented correctly. Most of the automatic analysis workflow(s) target identifying the animal, tracking its kinematics and/or body keypoints/pose from which behavior is derived [6]. Automatic workflows range from simple image analysis such as background subtraction or shape/color detection, which have mostly been applied in controlled lab settings, to complex machine learning algorithms that generalize well across different animals, behaviors, and environments [22, 14, 15].

Though machine learning has been demonstrated as a powerful tool to collect field animal behavior data [9], it usually relies on large amounts of accurate, manually-annotated ground truth datasets to train models, often leading to bottlenecks in analysis. Furthermore, it is desirable that the training datasets are generated in consultation with experienced researchers to ensure greater reliability in the model predictions. Therefore, exploiting the full potential of machine learning to study animal behavior in the wild, requires an easy-to-use tool that can assist in rapidly generating large amounts of high-quality annotated behavioral data while reducing the reliance on manual time-consuming effort.

Here, we present an open-source easy-to-use workflow, using zebras as an example, that can detect individuals in their natural environment and automatically infer their atomic scale behaviors of standing, grazing, walking, and running. Our workflow is based on the open-source annotation tool smarter-labelme which has previously been shown to reliably detect objects and animals [18]. We exploit its fast frame-by-frame object/animal detection capability, and develop on top of it a behavior classifier combined with a robust tracker of zebras. Our framework can thus detect and track all zebras in the scene and their behavior in a fast and reliable manner. In doing so, it presents a single tool that can be readily deployed to infer zebra behavior in the wild without expensive computational resources or in-depth technical know-how. Our workflow has the potential to be applied to other animal systems to rapidly generate high-quality large annotated datasets to be used for downstream tasks including training other networks or directly use for behavior analysis.

## 2. Methodology

### 2.1 Fast animal detection and behavior annotation workflow

Figure 2 describes the overall workflow of our proposed framework for fast behavior annotation, training, and deployment. It is developed on top of Smarter-labelme [18], and consists of three parallel, but inter-connected streams. The first stream (S1) focuses on detecting individual animals in the image frames, the second (S2) stream consists of manually classifying desired behaviors of the automatically tracked animals, and the third stream, (S3) focuses on training and deployment of the behavior inference classifier along with bootstrapping with S2.

**Figure 1:**
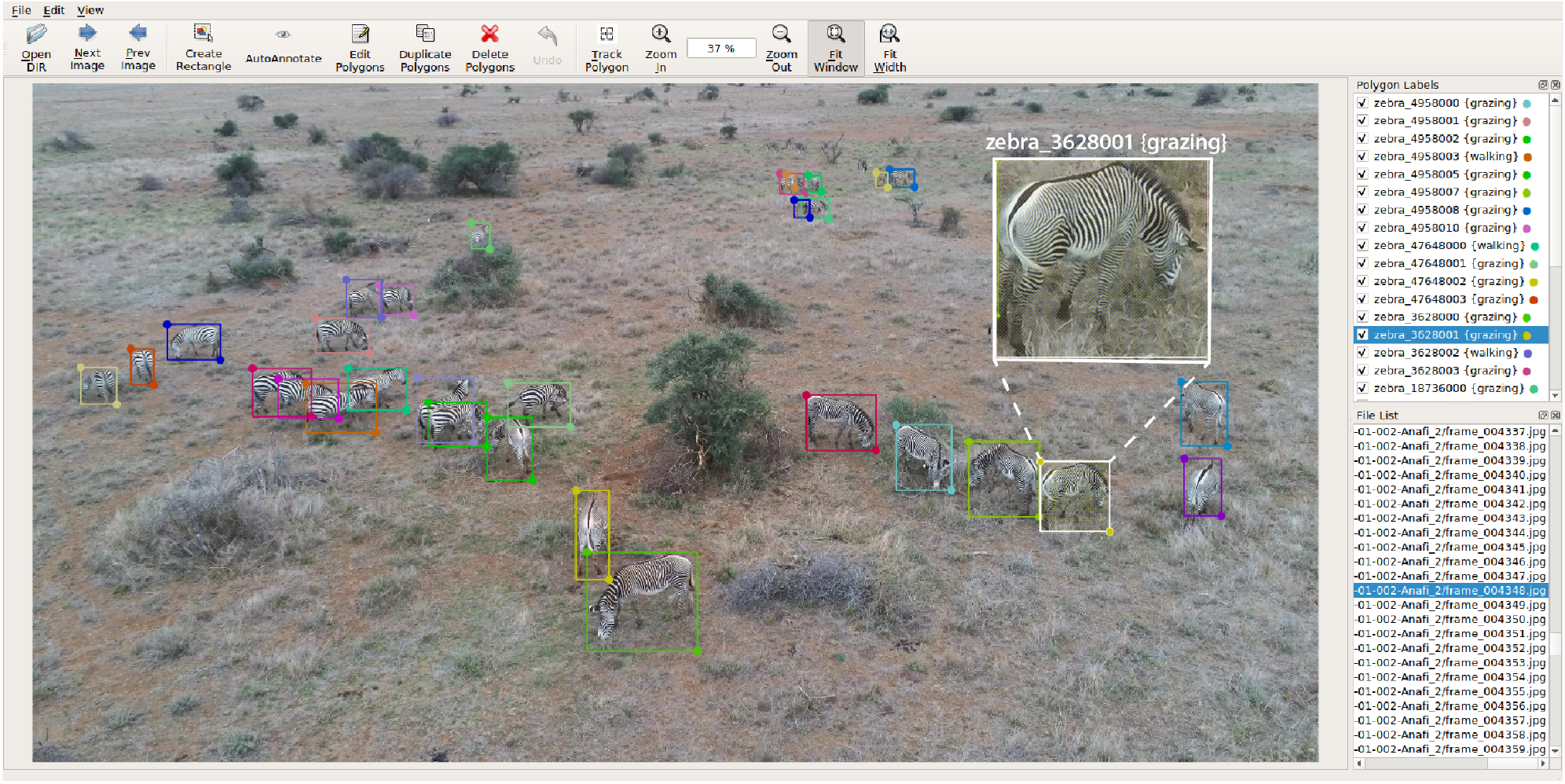
A snapshot of the smarter labelme interface showing the bounding box around each zebra. The ‘Polygon Labels’ panel displays the animal ‘id’ and the corresponding behavior detected/corrected. The bounding box colors correspond to the ‘ids’ of the animal. Inset shows the zoomed in view of one of the animals identified with its corresponding unique id and behavior.

**Figure 2:**
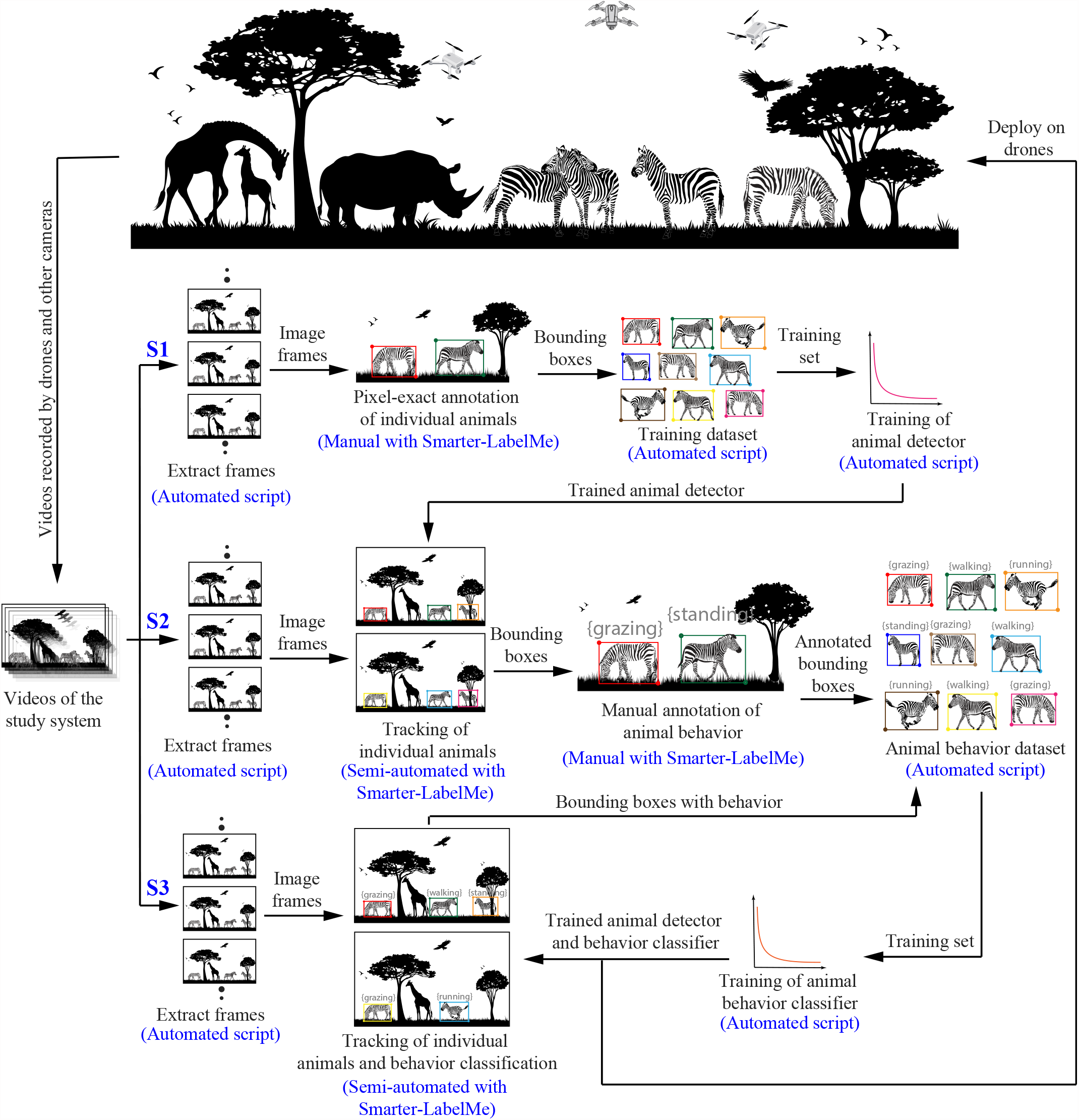
Architecture of the semi-automated animal detection and behavior inference workflow. The workflow is divided into three streams - S1, S2, and S3. The workflow is initiated from S1 by manually annotating pixel-exact bounding boxes of animals to generate animal detector training data followed by training the detector network [18]. The trained animal detector is used in S2 to perform manual annotation of behaviors and generate a training dataset to train the behavior classifier. S3 takes the output of S2 to semi-automate the entire behavior annotation process and rapidly generate more training data to fine-tune the behavior classifier and reach sufficient animal detection and behavior inference accuracy for deployment in field missions.

#### 2.1.1 Stream 1 (S1) - animal tracker

We assume to start with a large set of animal videos belonging to the study system. The objective of the first stream is to obtain enough pixel-exact annotations of the animals such that they capture sufficient variation within the context of the study and that the animal detector is trained with sufficient accuracy. This is usually achieved by selecting a small batch of videos, ensuring diversity in the time of day, light conditions, camera viewpoint, animal poses, and presence of multiple individuals. From these videos, image frames are extracted ensuring the aforementioned diversity. Extraction of frames from video can be easily achieved using the in-built frame extraction command of Smarter-labelme tool [18]. On these frames, manual annotation of pixel-perfect bounding boxes around the animal of interest are performed using the Smarter-labelme tool [18]. These annotations are then used to automatically generate a training set for the animal detector. The trained animal detector is the output of the first stream (See ‘S1’ in Figure 2), which is fed into the second stream (‘S2’ in Figure 2) for reasons as described further.

In our study, we achieved reliable zebra detections with 4283 annotations consisting of 34 unique zebras. These annotations were spread over 1067 frames extracted from 5 videos recorded over 2 days (Table 2, round 1). The videos were recorded using consumer-grade drones in the vicinity of the Mpala research center in Kenya.

#### 2.1.2 Stream 2 (S2) - semi-automatic behavior annotation

The workflow’s second stream objective (‘S2’ in Figure 2) is to employ a semi-automatic approach using Smarter-labelme to track the individual animals across frames. The tracking is achieved by fusing detections of animals using the trained detector from the first stream (S1), and a novel prediction method using RE^3^ [7]. The tracker produces bounding boxes and unique ids for each individual in the frame and tracks the individual’s identity across frames. The tracking of identity across frames is contingent on the frame rate at which images are extracted from the video. The frame rate should be such that Smarter-labelme’s tracking framework does not lose the animal across consecutive frames. The details of Smarter-labelme’s tracking mechanism along with considerations for consistent tracking are described in detail in Sec 2.3. After tracking the individual animals in a frame, the Smarter-labelme tool is used to manually annotate the desired behaviors for all individuals in the frame. The manual annotation is performed by individuals who are knowledgeable in the study system. The annotated behavior in a frame is carried forward to the next frame or backward to the previous frame, thereby requiring manual input only if the animal changes its behavior from one frame to the other. If bounding boxes exist for an individual across consecutive frames, group frame selection can be performed inside Smarter-labelme to change the behavior of the animal across all selected frames simultaneously.

An important distinction between S1 and S2 is that in S2 the bounding boxes produced by the animal detector are not required to be pixel-exact boxes. A more relaxed bounding box constraint further reduces the behavior annotation time while making the trained behavior classifier more robust to variations in bounding box positions with respect to the animal.

An automated script is then employed to generate training data from these behavior-annotated bounding box trajectories of the animals to train the behavior inference classifier, described in Sec 2.4. Both manual annotation steps mentioned above, for behavior inference classifier and animal detector, are essential for bootstrapping the auto-annotation capability of our proposed framework. This bootstrapping is further described in Sec. 2.1.3.

In our study, we annotated 158,516 instances of zebra behavior spread over 29,648 frames extracted from 19 videos recorded over the span of 5 days.

#### 2.1.3 Stream 3 (S3) - bootstrapping with minimal manual input

Stream 3 entails bootstrapping the entire process for fast animal tracking and behavior annotation to generate more training data from new videos for S2. Using the trained animal detector and behavior classifier from S2, Smarter-labelme tracks the bounding boxes and corresponding behaviors for all animals of interest in the frame.

A first pass of correcting all the bounding boxes and not the misclassified behaviors is performed. Once bounding boxes have been checked and/or corrected, the behavior of each animal is checked. Starting from the frame where a particular behavior starts, consecutive frames are browsed to check if the behavior is correctly classified. If a misclassification exists, all frames corresponding to the behavior are selected, and the behavior is simultaneously corrected by updating the behavior flag in the label field. Such an approach allows quick behavior annotation over large temporal periods, not requiring frame-by-frame correction for each individual.

The corrected behavior labels/classes are then added to the previous behavior classifier training dataset to expand the behavior classification training dataset further and quickly retrain the classifier for improved classification moving forward. These cyclic steps in S2 and S3 are performed until the behavior inference classifier achieves a sufficient level of accuracy. The trained behavior classifier from S2 can then also be deployed on downstream applications, such as on drones or other image/video recording equipment for real-time behavior inference in the field. Here, we do not provide any quantification of ‘sufficient’ accuracy since it is dependent on the study system.

### 2.2 Preliminaries and notations

Here, we introduce several notations that are used to describe the tracking and inference steps of our approach. Let **V** = {***i***_1_, ⋯, ***i***_*F*_} be a video, represented as a set of *F* consecutive image frames, where ***i***_*f*_ is the *f* ^th^ image frame. The corresponding annotation set is denoted as **A** = {***â***_1_, *⋯*, ***â***_*F*_}. Each image frame ***i***_*f*_ is a 2-dimensional pixel array of width *X* and height *Y*. Any (*x, y*)^th^ pixel of image ***i***_*f*_, denoted by ***p***_*x,y*_, is a 3 dimensional vector, 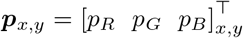 for all 0 *≤ x < X* and 0 *≤ y < Y* and *x, y* ∈ ℝ, where the components of the vector denote the red, green and blue (RGB) intensities of the pixel. [*x* = 0 *y* = 0]^T^ is assumed to be the upper left corner of the image frame. Each annotation vector ***â***_*f*_, corresponding to the image frame ***i***_*f*_, is a set, 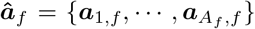, i.e., it consists of *A*_*f*_ individual annotations. Each annotation ***a***_*a,f*_ is itself a tuple, ***a***_*a,f*_ = {***l***_*a,f*_, ***b***_*a,f*_}, for all 0 *≤ a < A*_*f*_ and *a* ∈ ℝ, where ***l***_*a,f*_ is the label of the *a*^th^ annotation and ***b***_*a,f*_ the location vector of the *a*^th^ annotation. ***b***_*a,f*_ = [*x*_*a,f*_ *y*_*a,f*_ *X*_*a,f*_ *Y*_*a,f*_]^T^, where (*x*_*a,f*_, *y*_*a,f*_) are the pixel coordinates of the upper left corner of the rectangular bounding box (the annotation), and *X*_*b*_, *Y*_*b*_ are its width and height, respectively, in pixels. The label ***l***_*a,f*_ is a textual representation, which can be used to store information about the annotated object instance, such as its type, identity and behavior, along with other relevant metadata.

### 2.3 Semi-automated tracking of animal bounding boxes

In Smarter-labelme [18], on which our framework is developed, tracking is achieved by fusing detections of the objects with their predictions in each frame. While SSD multibox is used for detections [13], Re3 [7] is used to predict the annotated bounding boxes around those objects in subsequent frames. Re^3^, however, has the following shortcoming. When predicting objects in the subsequent frame, Re^3^ has a search area of exactly twice the annotated object in the previous frame, due to its network architecture. This typically produces correct predictions if only the object is moving and not the scene. However, if the camera itself is in motion and tilting or panning, which is usually the case with drone cameras, these predictions can quickly fail. Such camera motions can cause small and distant objects to shift, along with the whole visible field of view, by a multiple of the object’s size in pixels, even over a relatively short time-period. In this case, the new pixel coordinates of the object in the subsequent frame are out of the search area of Re^3^, causing an incorrect prediction which leads to tracking failure. It should also be noted that the annotator can also cause sudden shifts in the object’s pixel location/size/shape if the image frames are extracted at a low frame rate. For example, for a zebra running, images extracted at a low frame rate can cause sudden jumps in the individual’s location, causing the prediction to fail.

We address the camera motion-induced problem described above by first identifying a transformation matrix for the whole image, which describes the shift of the entire image in pixel-space. This correction matrix is then applied to the coordinates of the search area of each object before employing the tracker. To identify this transformation matrix, we use a parametric image alignment algorithm [4] on the down-sampled (to 100px × 100px) versions of the previous and the current image frames, ***i***_*f*−1_ and ***i***_*f*_, respectively. This has two benefits. First, although not GPU accelerated, the method is sufficiently fast on small images and needs to be executed only once for each pair of frames. Second, the influence of small moving objects in the scene on the algorithm is minimized by down-sampling, while the global image motion is maintained.

The tracking algorithm itself is then run as in [18], fusing Re^3^ [7] predictions with SSD multibox detections [13], both with shifted search areas. This dramatically improves the performance of the tracker on videos with moving cameras, such as drones cameras.

### 2.4 Behavior inference

We solve behavior inference as a classification task, using the Resnet34 [10] deep convolutional neural network (DCNN). Input to the network is a scaled and cropped region of image around the detected/annotated animal in each image frame. To obtain this, for every annotation, ***b***_*a,f*_, a square region of the image is selected, which is sized approximately *k* = 1.3 times the largest dimension of the annotation’s bounding box. Given ***b***_*a,f*_ = [*x*_*a,f*_ *y*_*a,f*_ *X*_*a,f*_ *Y*_*a,f*_] as the annotation, the largest dimension of the bounding box, denoted as *B*_*a,f*_, is given as

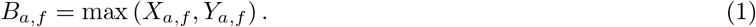

The scaled and cropped region of image, 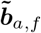, is then given as

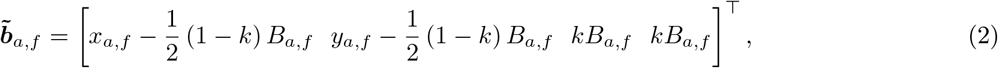

which is further rescaled to the network input size of 300px × 300px. The output classes are all relevant annotated behaviors, plus *unknown*. We integrate this network into smarter-labelme [18] to infer the behavior of the newly labeled or tracked animals, which allows the user to test the output of the network easily on new datasets, or use the network for assisted annotation. Labels with automatic behavior annotation are also marked as auto-labeled, to distinguish them from manually annotated labels.

#### 2.4.1 Training data generation

For training the behavior classifier, we select random crops of all behavior annotated animals while uniformly varying *k* between 1.25 and 1.67. The idea is to make the network invariant to the exactness of the bounding box detection. We balance the training data between behavior classes to prevent a network bias towards any specific behavior. Completely random crops of the annotated images are selected for the *unknown* class, which will, in most cases, not include a single animal. The motivation is to force the classifier to ensure that an animal is present for inferring the behavior and not rely entirely on background cues for the behavior classification.

#### 2.4.2 Training

At training time, random crops (*unknown* class) are fed to the network along with the corresponding ground truth annotations. The network weights are optimized through stochastic gradient descent based on a negative log-likelihood (nll) loss function. The training data is augmented at the start of training to randomly flip, blur, slightly crop, or slightly rotate the training image to increase variance. The color-space is also randomly adjusted to replicate aspects of different lighting conditions. These training data augmentation steps increase the diversity of data presented to the network for training and make the network generalize better. Altogether, our workflow results in a very light-weight training. In our experiments, shown later, the network converges in less than 1 hour of training time, over 12 epochs with 50000 training crops per epoch. We used an initial learning rate of 0.1 with a learning rate decay factor *γ* = 0.774 applied between epochs. These hyper-parameters were found by manual hyperparameter search comparing convergence speed and loss after a low number of epochs. We selected a batch size of 32 crops.

## 3. Datasets

In order to demonstrate our framework, we apply it on a video dataset of plains and Grévy’s zebras (*Equus quagga* and *Equus grevyi*). The dataset is split into training and test sets, in the ratio of 70 : 30. To make our experiments with the framework completely reproducible, we make the whole dataset available to the community^4^. In Table 1, we discuss the details of the collection procedure and the details of the data we make publicly available.

**Table 1:** Details of the dataset collection procedure and the publicly released data.

|  |  |
| --- | --- |
| Equipment | All videos in the dataset were recorded using four off-the-shelf drones: 2 DJI Mavic 2 Pro and 2 Parrot Anafi 4K. Videos from Mavics were recorded at 4K resolution (3840x2160 pixels at 29.97 fps). Videos from Anafis were also recorded at 4K resolution (4096x2160 pixels at 23.98 fps) |
| Location | Mpala, Kenya: A region characterized by arid and semi-arid savannahs and woodlands. |
| Date & Time | All videos were recorded during daytime (between 0800 and 1800 local time), in the months of July and August 2022. |
| Recording Procedure | The research team first searched for the presence of zebra herds at Mpala through manual observation from an off-road vehicle. After spotting the zebras, which typically happened during the first hour of the search, the vehicle was stopped approximately 200m from the herd. One or two drones were then manually deployed such that the take-off sound did not startle the herd; the drones were placed behind the vehicle, on the opposite side of the zebra herd. After take-off, the drones quickly ascended to an altitude of $\sim 100\text{m}$. From there, they were cruised, at a relatively constant altitude, towards the center of the herd, and then slowly descended up to 10–20m above the herd. This procedure was performed by the drone pilot by manually observing the drone and the zebra herd, simultaneously. Once the drones reached close to and above the zebras, they were flown following the zebras, keeping as many zebras as possible in their cameras’ field of view. To this end, the pitch angle of the camera gimbal and the yaw orientation of the drone were manually controlled by the pilot. While the drone cameras did not have any optical zoom, digital zoom was also never performed. Even during the period of approach (ascend and initial cruise), the pilot attempted to keep the animals in the drone camera’s field of view, however this was not always successful. The drones were kept following the herd until the battery levels approached a pre-determined threshold to return safely. At that point, the drones were manually flown back to the start position, near the vehicle. A complete flight was $\sim 25$ minutes, out of which the useful video recordings range from 5 to 20 minutes. |
| Released Dataset | • The extracted image frames from the video and corresponding annotations. Note that the dataset includes only the annotated frames, thus not all recorded video frames are part of the dataset.<br>• The generated dataset for the zebra detector training, merged with MSCOCO [12]<br>• The generated dataset (training, test1, test2) for classifier training and testing.<br>• The weights of the final trained zebra detector network.<br>• The weights of 10 trained behavior networks, each with a different seed. |

**Table 2:** Statistics of annotated data.

| Round | 1 | Round | 2 | Combined |  |
| --- | --- | --- | --- | --- | --- |
| Videos | 5 | Videos | 19 | Videos | 24 |
| Frames | 979 | Frames | 29648 | Frames | 30627 |
| Zebras | 34 | Zebras | 224 | Zebras | 258 |
| Annotators | 4 | Annotators | 9 | Annotators | - |
| Annotation Time (hours) | 27 | Annotation Time (hours) | 112.25 | Annotation Time | - |
| Annotations Total | 4283 | Annotations Total | 158516 | Annotations Total | 162799 |
| Of those: grazing | 1268 | Of those: grazing | 70466 | Of those: grazing | 71734 |
| standing | 824 | standing | 33072 | standing | 33896 |
| walking | 1848 | walking | 41387 | walking | 43235 |
| running | 343 | running | 13591 | running | 13934 |

## 4. Experiments and Results

### 4.1 Implementation

We begin with describing the manual annotations performed in stream ‘S1’ of our methodology (see Figure 2) for training the zebra detector. Here we instruct the annotators to make pixel-exact annotations of all individuals. To maximize the diversity in this data, we extract videos at a low framerate (1 Hz) to reduce similarity in subsequent frames while still allowing the tracker to accelerate the annotation across frames. A total of 4283 bounding boxes were annotated across 979 annotated video frames (see Table 3). From this data, we trained SSD Multibox [13] as the animal detector. To do so, we extracted 21000 random crops from those 979 frames, such that they contained one or more annotated bounding boxes of the animal, combined it with the 118,287 annotated frames from the MS-COCO 2017 training data [12] and then used all of that to train the detector. Doing so took *∼* 24 hours on an NVIDIA GeForce RTX 2080 Ti GPU. The resulting detector showed comparable performance to pretrained networks on the MS-COCO dataset, but was optimized for our data distribution. No new classes had to be added since zebra is already a pre-defined class in MS-COCO, however with only 5304 individual annotations across different zebra species, in different habitats, and typically in closeups. By adding an additional 21000 annotations, all matching our dataset distribution, the detector became sufficiently accurate to detect, with high accuracy, most zebras in all video frames.

**Table 3:** Training, and test sets for behaviour detector training.

| Set | Train | Set | Test set 1 | Set | Test set 2 |
| --- | --- | --- | --- | --- | --- |
| Crops per class | 10000 | Crops per class | 1000 | Crops per class | 2000 |
| Zebras | 179 | Zebras | 27 | Zebras | 52 |
| grazing | 51056 | grazing | 6115 | grazing | 14545 |
| standing | 28600 | standing | 1501 | standing | 3784 |
| walking | 33516 | walking | 1605 | walking | 8107 |
| running | 8062 | running | 1717 | running | 4025 |

Next, for the stream ‘S2’, we annotated 29648 frames in 19 videos for a total of 158516 zebra annotations. In this case, frames were extracted at 8 FPS to ensure that the automated tracker was predominantly successful (see Section 2.3) and the annotators could mostly rely on automated tracking and focus on behavior annotation. The final behavior classifier training dataset consisted of the 158516 behavior annotations along with the annotations from ‘S1’ to which behavior information was added. Annotators were instructed to distinguish between 4 easily identifiable individual behaviors. “grazing”, implying the animal was eating grass or leaves; “standing”, implying the animal was stationary but not eating, ‘standing’ also included other activities such as self-grooming, sleeping, or vigilant; “walking”, implying the animal was moving at a leisurely pace, while not eating; “running”, implying the animal was moving fast. This led to a combined behavior-annotated dataset of 71734 grazing, 33896 standing, 43235 walking, and 13934 running zebra annotations. (Table 2)

From the total of 30627 annotated frames (from ‘S1’ and ‘S2’ combined) we created a training set of 50000 training images for the behavior classifier. In that, 10000 random crops were labeled “unknown” – these typically contain background or multiple cropped animals, while the rest 40000 were zebra instances with their behavior labels, 10000 per behavior label. These were cropped out from the original images between 1.25 and 1.67 times the size of the zebra bounding box. We also created 2 test sets. Test set 1 was with 5,000 total images, 1000 per class, and Test set 2 with 10000 total images, 2000 per class. For this, the total zebra ids of 258 in the original dataset were split. 179 zebra ids were assigned as training instances, 27 zebra ids were assigned to Test set 1, and 52 zebra ids to Test set 2 in a 70/10/20 data split.(Table 3)

We trained Resnet34 [10] on the train set, using stochastic gradient descent with a learning rate decay of *γ* = 0.774 after every epoch. Good convergence was reached after 12 epochs which took *∼* 1 hour. The short training time allowed manual hyperparameter tuning to find acceptable initial learning rate and decay values, which can be found as default values in the provided training code. We trained 10 networks, using different random seeds, and then compared their performance on both validation and test set. The random seed affects both the random initial weights of the network, the order of training samples, and the applied random train time data augmentation. Therefore, no two networks are exactly the same, and depending on these random numbers can achieve slightly better or worse performance, both on individual classes and overall. In cases where classes are hard to tell apart, this can also affect the network’s overall bias favoring one class over another. This is visualized in Figure 7.

### 4.2 Accuracy of Behavior Inference

Table 4 and Figure 7 show the overall accuracy of 10 networks on two different test sets – Test set 1 and 2. Each of the 10 networks are trained with a different seed. On Test set 1 we reach a maximum accuracy of 78% and on Test set 2 we reach 81%. The accuracy on Test set 1 and 2 demonstrates the reliability of the network to identify the 4 behaviors and also acts as an indicator of the performance on previously unseen data. We reach classification accuracy of well over chance, which would be 25% for 4 classes presented in equal amounts in the test sets. Figure 3 and 4 show the per class performance on both test sets as well as the percentage of mis-identifications per class. Except for 3 networks on Test set 1 (Figure 3), all networks reached a per-class accuracy of over 50% for all classes.

**Table 4:** Accuracy of behaviour classifier on both test sets for 10 trained networks.

| Network | Test set 1 | Test set 2 |
| --- | --- | --- |
| 0 | 0.76 | 0.81 |
| 1 | 0.75 | 0.80 |
| 2 | 0.74 | 0.79 |
| 3 | 0.76 | 0.80 |
| 4 | 0.77 | 0.80 |
| 5 | 0.77 | 0.80 |
| 6 | 0.75 | 0.79 |
| 7 | 0.77 | <u>0.81</u> |
| 8 | 0.77 | 0.78 |
| 9 | <u>0.78</u> | 0.79 |

**Figure 3:**
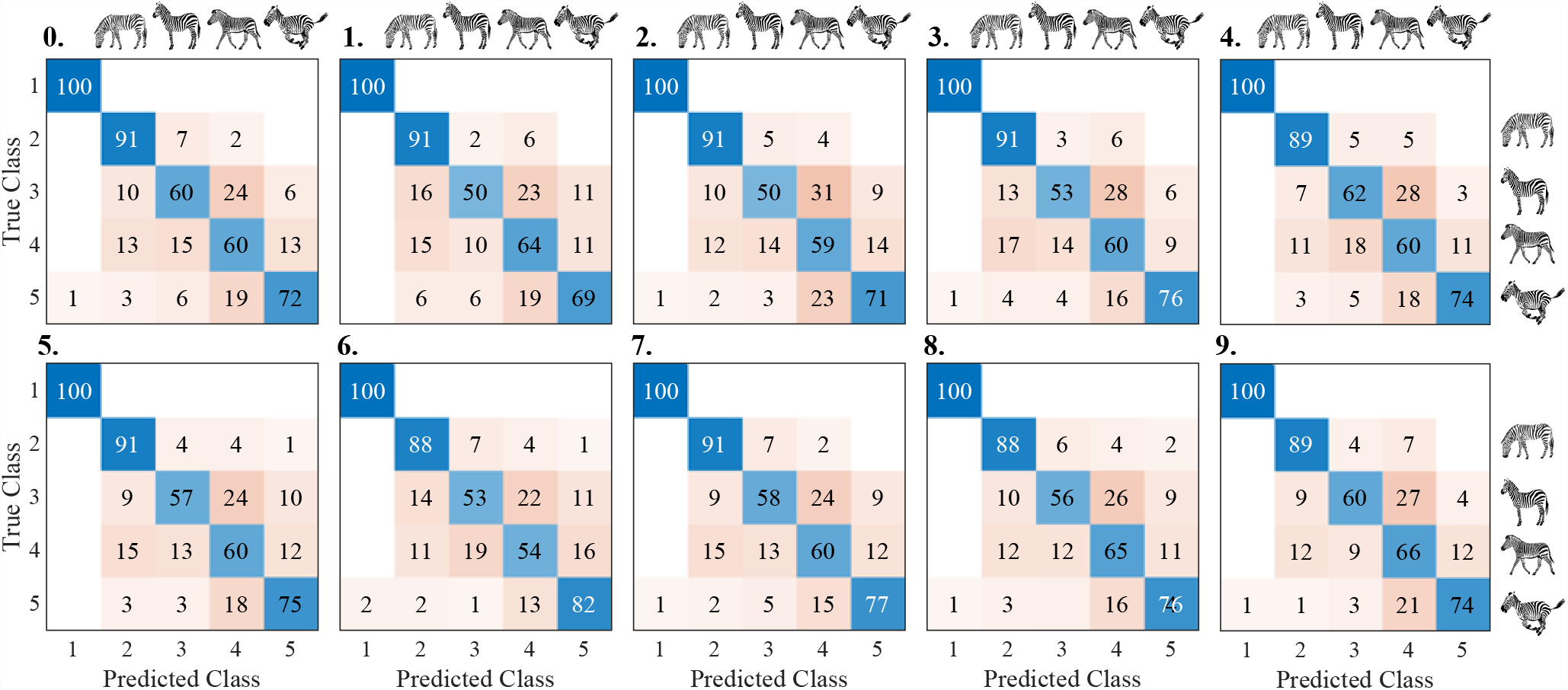
Groundtruth vs. Detection Confusion matrix for 10 trained networks - Test set 1. Each row shows the percentage of detections (per column) for each ground truth class in the order: “unknown”, “grazing”, “standing”, “walking”, “running”.

**Figure 4:**
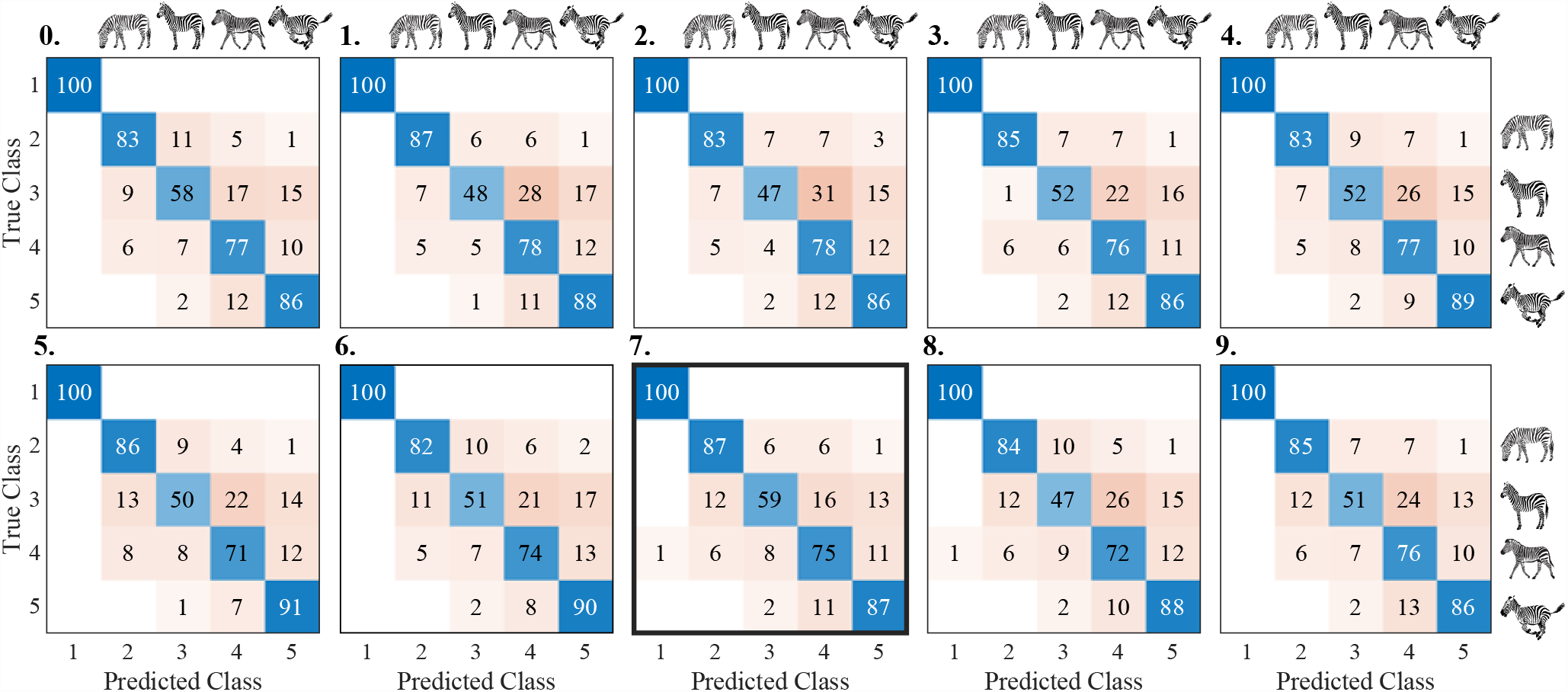
Groundtruth vs. Detection Confusion matrix for 10 trained networks - Test set 2. Each row shows the percentage of detections (per column) for each ground truth class in the order: “unknown”, “grazing”, “standing”, “walking”, “running”.. The confusion matrix for the best performing network (network 7) is highlighted with a bold boundary.

All networks learned to identify the presence of an animal in the picture, leading to negligible confusion between the “unknown” class and all other classes. “Grazing” and “running” zebras were also reliably distinguished, with almost no confusion between these cases. The confusion between “grazing” and “standing” zebras was slightly higher, similar to the confusion between “grazing” and “walking”.

We observed a moderate confusion between “standing” and “running” as well as “walking” and “running”, and the highest confusion between “standing” and “walking”. In the latter case, consistently more “standing” zebras were misidentified as “walking” than the other way around.

Figures 5 and 6 provide plausible explanations for the achieved accuracy. Shown in these figures are the class activation matrix (CAM) for both correctly and incorrectly classified examples from Test set 2. It indicates which spacial regions of the image contributed, and to what extent, towards the classification task. In other words, where the network was “looking at”. In figure 5, we show two examples each for correctly identified behaviors.

**Figure 5:**
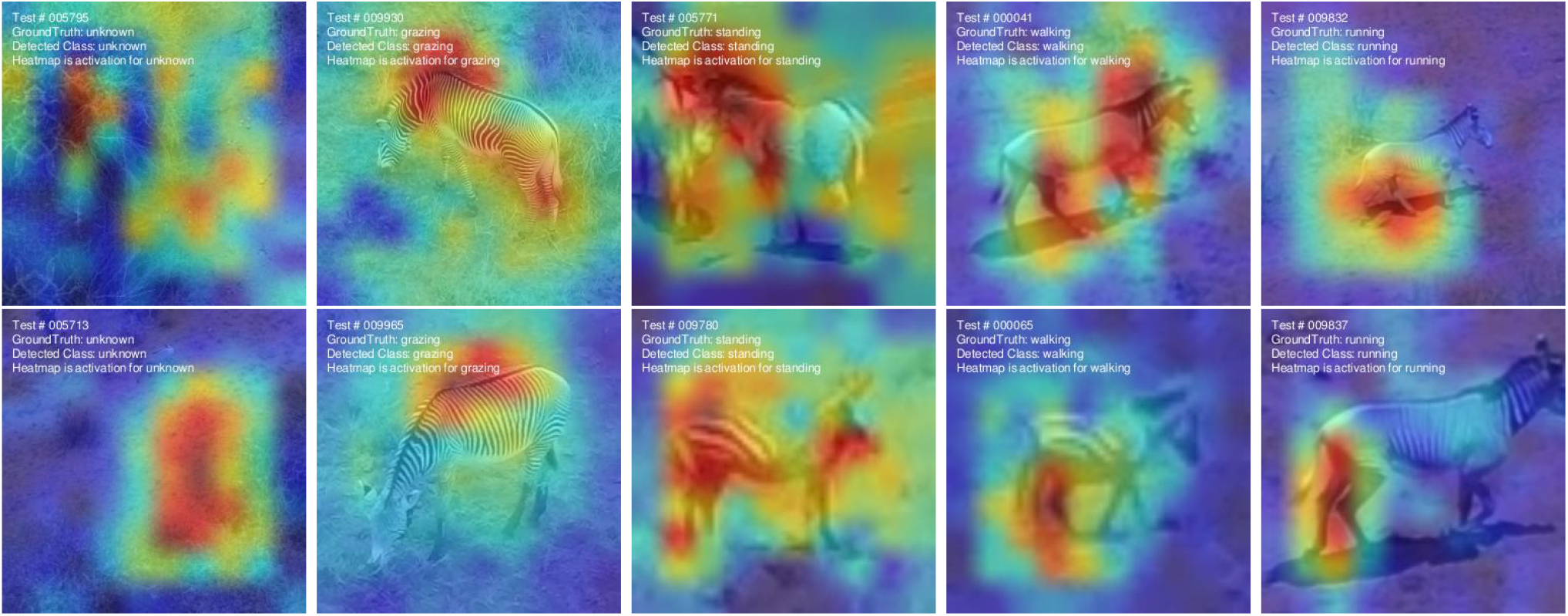
Class Activation Heatmaps (CAM) for successful examples on the Test set 2 with network 7. The spacial attention of the network suggests that it uses the absence of a raised head as an indicator that the zebra is grazing. To distinguish between standing, walking, and running, the network learned to pay attention to the legs and likely identifies the behavior class based on relative leg stance.

**Figure 6:**
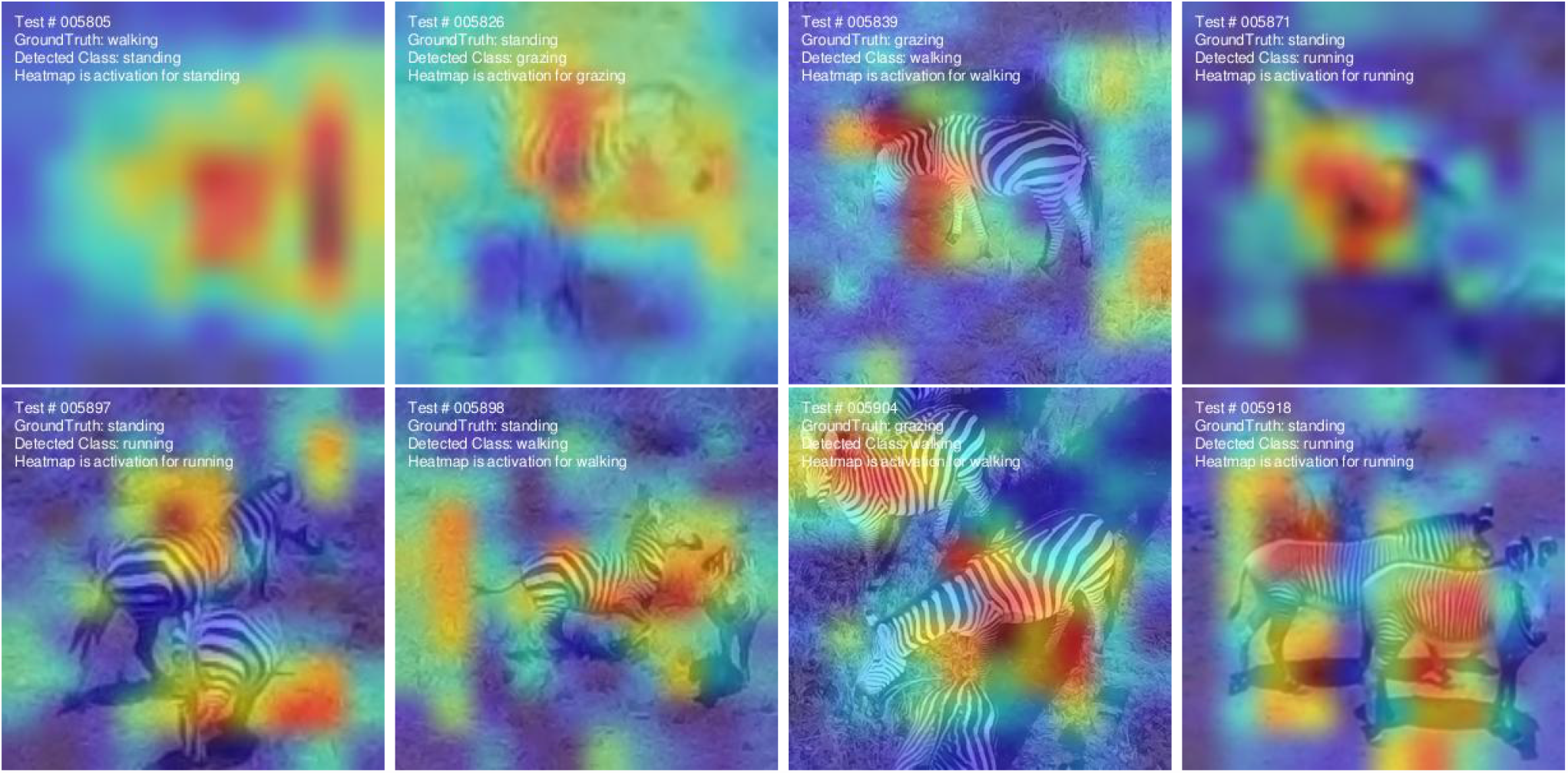
Class Activation Heatmaps (CAM) for failure cases. Prominent failures happen when the zebra is very distant and hard to distinguish, but also when there are multiple zebras in close proximity, confusing the classifier with conflicting information. Steep viewing angles and simultaneous actions (grazing while walking) also seem to sometimes confuse the network.

**Figure 7:**
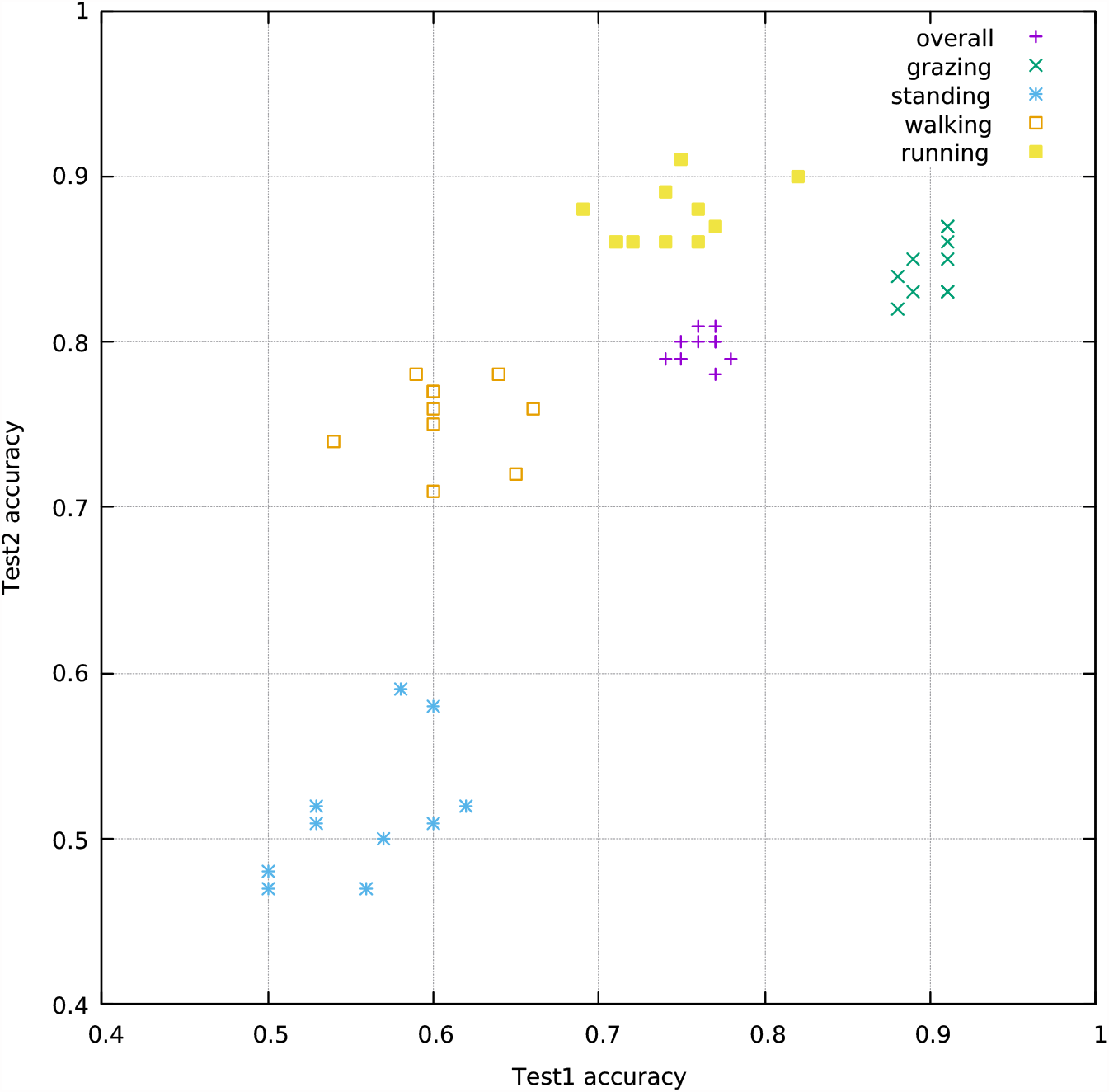
Cross-correlation of network performance on Test set 1 and Test set 2 for the whole dataset as well as for each behavior class.

The “grazing” class shows visible activation on the zebra body with a maximum where the head is not present but would be if the zebra was upright. This seems to allow the network to identify a grazing zebra by the lowered head, even if the head is not visible. This is especially advantageous when the zebra is observed from the posterior end and the head is occluded by the body.

The “standing” class shows multiple activation areas across the zebra body, both on the head and torso as well as the legs, while the “walking” and “running” classes primarily seem to focus on the legs of the animal for distinction. This provides a plausible explanation for the confusion between these classes. A zebra with all 4 legs straight and head up is almost certainly standing while a zebra with legs in a stride stance is more likely to be in motion, but it is possible for a zebra to stand still in this stance. The network identifies these instances as “walking”, which explains the asymmetric confusion. Much more “standing” zebras are misidentified as “walking” than “walking” zebras are misidentified as “standing”.

The confusion between “walking” and “running” is more symmetric in nature. The only distinguishing factor the network highlights is the articulation of the legs, where some articulations are highly indicative for running there remains some overlap with “walking” depending on the gait.

Another factor, which is apparent in the failure cases in figure 6 is that the zebra, or its distinctive features might not be clearly visible due to blur, occlusion, or the presence of a second animal in the frame. Aside from conflicting features provided by a second animal, interactions between zebras also lead to dynamic poses which are otherwise rare and not correctly interpreted by the network or simply fall outside of a simple 4-class behavior set.

Both, the possible poses of zebras as well as zebra behavior in itself form a long-tailed distribution, with some prominent behaviors forming the vast majority of available data, while certain interactions are very rare.

Figure 7 shows the cross-correlation between the accuracy on each test set (Test set 1 and 2) for the overall network as well as for each individual class (except “unknown”). The wider spread for the hard to distinguish classes, in combination with the confusion matrices, suggests that each network has a different prior, trading accuracy for one class against another, with the same overall accuracy, and same loss at train time. This could be influenced by deliberately providing an imbalanced training set, which would cause the network to prefer the more prominent class. A weak positive correlation between the accuracy of Test set 1 and Test set 2 is visible for most classes. This suggests that choosing network from an ensemble that is performing well on a known test set is a good strategy to also perform well on unknown data.

## 5. Outlook

### 5.1 Atomic behavior to long-term behavior

Identifying atomic behaviors in each video frame allows numerous possibilities to characterize and understand behaviors over a longer period of time (e.g., activity time budget) as well as extend them to infer higher-level behaviors of the animal and the group (e.g., hunting, conspecific interactions). Figure 8 demonstrates a simple example of extending the atomic behaviors that are classified in our zebra dataset to infer the activity budget of each individual. Based on the study system and the requirement, the Smarter-labelme behavior classifier can be trained to classify any range of behaviors. For example, in the case of zebras, the range of behaviors classified can be extended to include instances of ‘self-grooming’, ‘urinating’, ‘drinking’, ‘defecating’, ‘mutual grooming’, ‘mating’, and so on - providing unprecedented levels of detail about the animal’s and the group’s activity.

**Figure 8:**
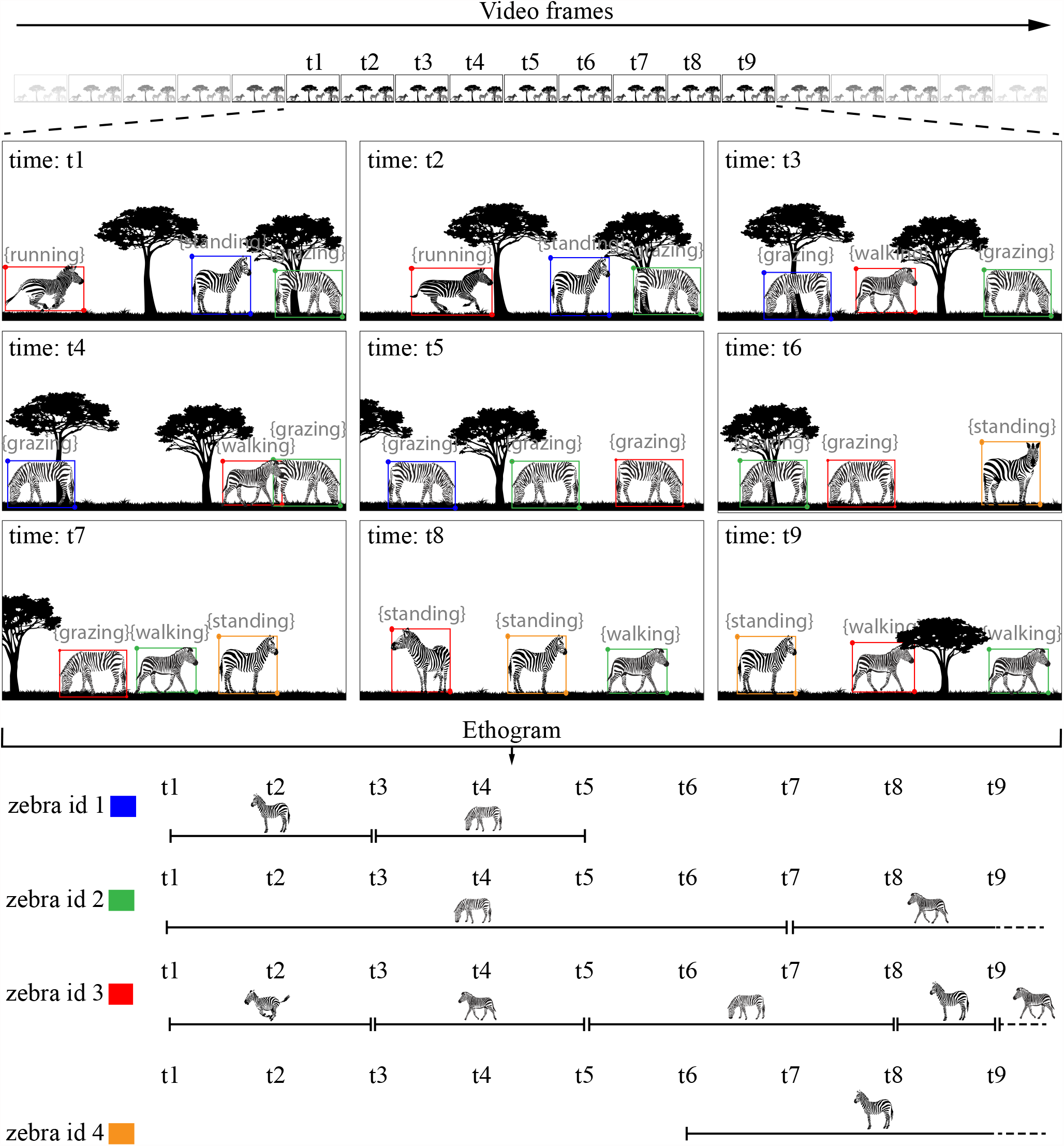
An illustration showing how atomic scale behaviors measured from our workflow can be scaled to long-term behaviors of the animals in their natural habitat. The unique colors correspond to each of the individuals tracked over a temporally aligned sample time interval from ‘t1’ to ‘t9’. The temporal resolution of each of the inferred long-term behaviors will depend on the time difference between each of the sampling intervals.

Given the confidence we have in identifying the behavior of individuals we can then use this approach to identify differences among individuals of differing species or even of different sexes or reproductive states within species. For example, in the videos shown above both plains and Grevy’s zebras were in a mixed species herd. When together, do they act more similarly in terms of standing as they look for predators or reproductive competitors? Or are there species differences that might persist irrespective of the nature of the herd they are in. Similarly, within a species, can males be identified by their behavioral phenotype from females? If possible this would be extremely valuable since many species which are not sexually dimorphic in size or armaments are hard to sex. And even within a sex, it might be possible to assess a female’s reproductive state since lactating females may adopt behaviors at different relative frequencies from non-lactating females that don’t have to cater to offspring.

### 5.2 Rapidness and advantage of the annotation framework

The most common bottleneck of machine learning approaches in the field of animal behavior is the reliable annotation of large amounts of datasets to train machine learning models. Our framework addresses this bottleneck by providing a semi-automated workflow to produce large and reliable annotated datasets without requiring significant manual efforts. This allows the rapid evolution of trained networks to include more behaviors, increase their accuracy, and the possibility of deploying in real-time in the animal’s natural habitat. Based on the study system and requirement, the annotated datasets can also be used directly for behavioral analysis.

In our study, the annotation time was reduced from 22.69 s to 2.55 s per annotation per individual (see Table 2).Though the reported annotation time is subjective and depends on a multitude of factors, including the annotator’s previous experience with the animal/workflow, it provides a promising insight into the potential future speed benefits gained from our workflow while not compromising the quality of the produced dataset. Furthermore, the semi-automated workflow allows animal behavior experts to quickly annotate diverse behaviors thus reducing the possibility of errors that can arise through annotations performed by individuals who are not familiar with the study system (e.g., volunteers, outsourcing annotation tasks to online platforms).

Overall, a method to quickly and accurately infer behavior in every video frame can revolutionize the field of animal behavior and unlocks the potential to study behavior at large spatiotemporal scales in the animal’s natural habitat. A key bottleneck of such a behavior inference method is obtaining large sets of behavior-annotated video data. Our approach not only addresses this bottleneck but also presents a method for behavior inference, using that approach. We demonstrate our overall framework on four behaviors (standing, walking, running, and grazing) of plains and Grévy’s zebras using a light-weight network that can be trained in approximately 1 hour. The relatively fast training and inference speed of the network makes it an attractive candidate for future deployment to infer real-time behavior classification in the field. While we reach high accuracy for grazing and running, we discuss the potential reasons for walking and standing behaviors to have comparatively lower accuracy, which should provide pointers to tackle this issue in the future. We further discuss the future directions in which our approach will be useful to answer a variety of challenging questions in animal behavior. Lastly, we provide the complete code and dataset for the research community for easy adoption in other machine learning workflows and animal behavior studies.

## Supporting information

Video Abstract

## Footnotes

1 Code: https://github.com/robot-perception-group/animal-behaviour-inference

2 Data: https://keeper.mpdl.mpg.de/d/a9822e000aff4b5391e1/

3 Video Abstract: https://youtu.be/Zu-t0JJsz5o

4 Drone Video Dataset of Grévy’s and Plains Zebras: https://keeper.mpdl.mpg.de/d/a9822e000aff4b5391e1/

